# Carafe2 enables high quality *in silico* spectral library generation for timsTOF data-independent acquisition proteomics

**DOI:** 10.64898/2026.03.27.714846

**Authors:** Bo Wen, J. Sebastian Paez, Chris Hsu, Daniele Canzani, Alexis Chang, Nicholas Shulman, Brendan X. MacLean, Matthew D. Berg, Judit Villén, William E. Fondrie, Lindsay K. Pino, Michael J. MacCoss, William S. Noble

**Affiliations:** Department of Genome Sciences, University of Washington; Talus Bioscience, Inc. Seattle, WA, USA; Paul G. Allen School of Computer Science and Engineering, University of Washington

## Abstract

Data-independent acquisition (DIA) proteomics enables reproducible and systematic peptide detection and quantification, and trapped ion mobility spectrometry (TIMS) on the timsTOF platform further improves DIA by synchronizing ion mobility separation with quadrupole precursor sampling. Analyzing the highly multiplexed spectra generated by DIA typically relies on spectral libraries, and fully leveraging the additional ion mobility dimension requires these libraries to include accurate retention time, fragment ion intensity, and ion mobility annotations. Existing *in silico* spectral library generation tools either lack ion mobility support entirely or rely on models trained on data-dependent acquisition (DDA) data, that can introduce a mismatch that may not capture unique experiment-specific biases when applied to each respective timsTOF dataset. Carafe is a software tool that uses deep learning models to generate high-quality, experiment-specific *in silico* libraries by training directly on DIA data. In this study, we extend Carafe to generate libraries for timsTOF DIA data, which involves fine-tuning retention time (RT), fragment ion intensity, and ion mobility prediction models using timsTOF DIA data. Carafe2 operates directly on native timsTOF raw data (Bruker .d directories) without the need for data conversion. We demonstrate the performance of Carafe2 across a wide range of DIA applications, including global proteome, phosphoproteome, and plasma proteome datasets. Comparing Carafe2 fine-tuned RT, fragment ion intensity, and ion mobility prediction models with pretrained DDA models, we find that Carafe2 models outperform pretrained models on a variety of DIA datasets. We then demonstrate the utility of *in silico* libraries generated by Carafe2 for peptide detection on several different types of timsTOF DIA datasets by comparing with the libraries generated with DDA-trained AlphaPeptDeep models, DIA-NN built-in models, and empirical spectral libraries generated from DDA experiments.

## 1 Introduction

Mass spectrometry-based proteomics has become an indispensable tool for studying complex biological systems [1, 2]. The vast majority of mass spectrometry proteomics experiments are performed using a bottom-up approach, in which proteins are extracted from biological samples and digested into peptides using a specific protease, typically trypsin. These peptides are then separated by liquid chromatography (LC) and introduced into the mass spectrometer, where precursor ions are selected and fragmented to produce tandem mass spectra (MS/MS) that serve as digital fingerprints for peptide detection.

Data-dependent acquisition (DDA) has been the primary acquisition strategy for bottom-up proteomics and is particularly powerful for detecting peptides in complex mixtures. However, because DDA relies on stochastic sampling — selecting precursor ions for fragmentation in real time based on their intensity — this approach suffers from poor sampling reproducibility across runs and incomplete coverage of lower-abundance species, making reproducible peptide quantification across large sample cohorts challenging without the use of strategies such as tandem mass tags [3].

Data-independent acquisition (DIA) addresses this sampling limitation by systematically fragmenting all precursor ions within predefined *m/z* isolation windows, thereby acquiring fragment ion data for every detectable peptide in each run [4–6]. In DIA, extracted precursor-to-product ion chromatograms (XICs) can be used in a manner analogous to targeted parallel reaction monitoring [7], enabling reproducible and comprehensive peptide detection and quantification across experiments. Although DIA is a promising strategy, the sampling of different precursor ions using a quadrupole mass filter remains somewhat inefficient. Only a single *m/z* window can be sampled at a time and the isolation windows must be wide enough to efficiently cycle through the *m/z* space in a chromatographic timescale. To overcome this sampling inefficiency, trapped ion mobility spectrometry (TIMS) has been implemented to improve the sampling of the ion beam prior to the quadrupole mass filter [8]. In this design, ions are first accumulated in an ion mobility trap and then released in order of their collisional cross-section (CCS), which is correlated with *m/z*. This correlation enables the quadrupole to synchronize its isolation window with the ion mobility separation, dramatically improving the efficiency of precursor sampling in a strategy called parallel accumulation-serial fragmentation combined with data-independent acquisition (diaPASEF) [9]. While diaPASEF greatly improves the utilization of the ion beam from the electrospray source, the ion mobility separation itself provides an additional dimension of information — the collisional cross-section — which is represented as a drift time and, while broadly correlated with peptide mass and charge, remains challenging to predict accurately at the peptide-specific level from sequence alone.

Because DIA fragments multiple precursor ions within each isolation window, peptide detection and quantification typically rely on spectral libraries that encode the expected properties of candidate peptides. Recent deep learning models have enabled accurate prediction of retention time and fragment ion intensities, making high-quality *in silico* spectral libraries a practical alternative to empirically generated libraries [10– 14]. By providing an additional dimension of separation, ion mobility enables peptide detection tools to better distinguish target peptides from co-eluting interferences with similar *m/z* [15]. Fully leveraging this dimension, however, requires libraries that include accurate ion mobility annotations alongside retention time and fragment ion intensity predictions. We and others have previously demonstrated that *in silico* spectral libraries generated using deep learning models can achieve superior performance compared to empirical libraries derived from DDA data or DIA data [11, 12, 14]. However, existing *in silico* spectral library generation tools primarily rely on models trained on DDA data [11, 13, 16, 17], which may not optimally capture the characteristics of DIA data, particularly from timsTOF instruments. In addition, although deep learning has enabled significant improvements in the accuracy of ion mobility prediction [13, 15, 18–20], most previously published *in silico* spectral library generation tools do not support ion mobility prediction [11, 14, 16, 21, 22]. To address the DDA–DIA mismatch in predicted libraries, we previously developed Carafe [14], a software tool that generates high-quality experiment-specific *in silico* spectral libraries by training deep learning models directly on DIA data.

In this study, we extend Carafe to support the generation of *in silico* libraries specifically tailored for timsTOF DIA data by fine-tuning retention time, fragment ion intensity, and ion mobility prediction models using timsTOF DIA datasets. To reduce preprocessing overhead and enable efficient access to the additional ion mobility dimension, Carafe2 operates directly on native timsTOF raw data (Bruker .d directories) via a standalone Rust-based raw-data access tool, TimsQuery, which was developed in this study, eliminating the need for conversion to intermediate formats such as mzML. Because the additional ion mobility dimension increases the size and hierarchical structure of timsTOF DIA files, efficient access typically requires a dedicated vendor software developer kit or an equivalent programming library; TimsQuery provides a lightweight alternative that meets this requirement. To make Carafe2 broadly accessible, we also provide a graphical user interface (GUI) that supports three common workflows. Finally, to facilitate the inspection of timsTOF DIA data analyzed with *in silico* spectral libraries generated by Carafe2, we also developed Timsviewer, a standalone visualization tool. We demonstrate the performance of Carafe2 on a variety of timsTOF DIA datasets, showcasing its ability to improve peptide detection and quantification compared to libraries generated using DDA-trained models, built-in models in DIA-NN, and empirical spectral libraries generated from DDA experiments.

## 2 Results

### 2.1 An overview of Carafe2

Carafe2 is an extension of Carafe [14] that enables the generation of experiment-specific, high-quality *in silico* spectral libraries specifically tailored for timsTOF DIA data. As illustrated in Figure 1, Carafe2 follows a systematic workflow to fine-tune deep learning models for predicting retention time (RT), fragment ion intensity, and ion mobility (IM) using timsTOF DIA datasets. Throughout this manuscript, we use “ion mobility” to refer to the instrument-reported inverse reduced ion mobility 1/*K*_0_ values, while noting that these values can be converted to CCS. The process begins with the generation of training data from a selected timsTOF DIA run, where peptides are detected either using DIA-NN in library-free mode or using other peptide detection tools capable of analyzing timsTOF DIA data.

**Figure 1:**
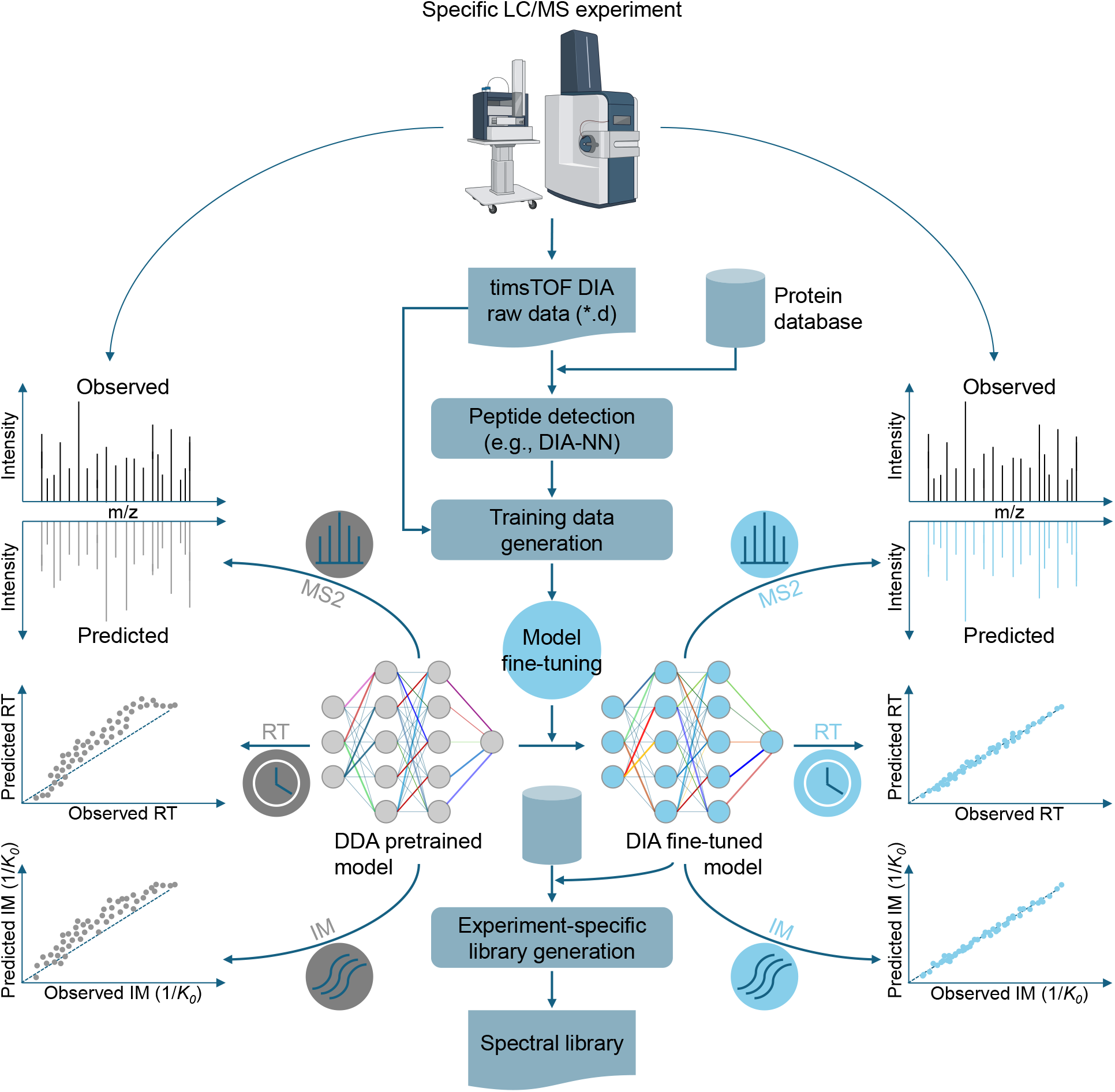
Overview of Carafe2. Carafe2 operates directly on native timsTOF raw DIA data together with peptide detections to fine-tune DDA data pretrained retention time (RT), fragment ion intensity (noted as “MS2”), and ion mobility prediction models, enabling generation of experiment-specific *in silico* spectral libraries. The ion mobility value is reported as the inverse reduced ion mobility (1*/K*_0_).

After peptide detection on a single or multiple DIA MS runs generated from a specific experiment condition of interest, Carafe2 uses the standalone Rust-based tool TimsQuery to extract chromatographic and spectral data directly from the native raw files across retention time and ion mobility, constructing experiment-specific training examples without file conversion. Carafe2 then fine-tunes the RT, fragment ion intensity, and IM prediction models, which are subsequently used to generate experiment-specific *in silico* spectral libraries that can be used by DIA-NN [23], Skyline [24], and FragPipe [25].

To make Carafe2 broadly accessible, we developed a graphical user interface (GUI) that provides an intuitive way to configure and run Carafe2 workflows (Supplementary Figure 1). The GUI supports three common workflows: (1) spectral library generation starting from an existing DIA-NN search result, (2) spectral library generation with DIA-NN search performed within the same workflow, and (3) an end-to-end DIA analysis in which Carafe2-generated libraries are used together with DIA-NN for peptide detection and quantification. To further improve accessibility for the user community, we have also integrated Carafe2 into the widely used Skyline tool [24].

### 2.2 Fine-tuning improves fragment ion intensity, RT and IM predictions on diverse timsTOF DIA datasets

To evaluate the accuracy of Carafe2’s fragment ion intensity predictions on timsTOF DIA data, following the protocol adopted in our previous work [14], we compared the performance of Carafe2 fine-tuned models with the pretrained DDA model from AlphaPeptDeep [13], thereby testing whether the DIA fine-tuning step in Carafe2 is beneficial. For this analysis, we used three timsTOF DIA datasets generated on two timsTOF MS instruments: two global proteome datasets and one phosphoproteome dataset. As shown in Figure 2a, each dataset contains DIA data generated from two different types of samples, a human sample and a yeast sample. For each dataset, we used a single DIA run (global proteome datasets) or multiple DIA runs (the phosphoproteome dataset, three replicates) from human as training data and a single DIA run from yeast as testing data. To evaluate the performance of the fine-tuned models, we used the Spearman correlation between the observed and predicted fragment ion intensities. Carafe2 masks peaks that are deemed likely to suffer from interference (see “Methods”), so only the unmasked peaks are used for evaluation.

**Figure 2:**
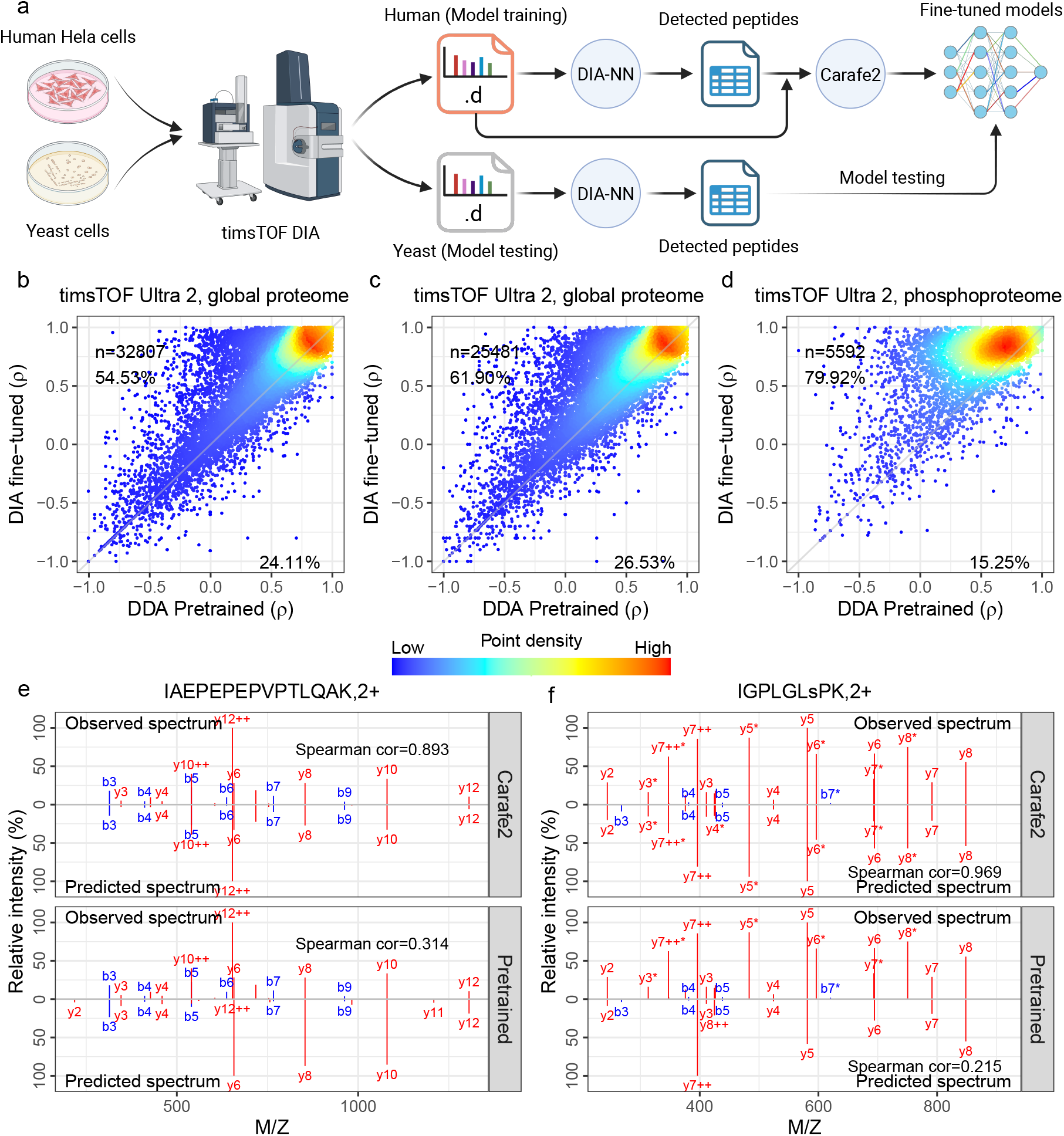
Fragment ion intensity model fine-tuning evaluation. **(a)** Overview of model evaluation scheme using paired human and yeast DIA datasets. **(b–d)** Fragment ion intensity prediction performance, measured by Spearman correlation (*ρ*), comparing the pretrained DDA model from AlphaPeptDeep (x-axis) with DIA fine-tuned models (y-axis) across three datasets. **(e–f)** Two examples of fragment ion intensity predictions using Carafe2 fine-tuned models and an AlphaPeptDeep pretrained model. The first peptide IAEPEPEPVPTLQAK with precursor charge state 2+ was detected in the dataset for (b), and the second phosphopeptide IGPLGLsPK with phosphorylation at S7 was detected in the dataset for (d). Panel **(a)** was created using BioRender.

The results of this evaluation show the benefits of fine-tuning with timsTOF DIA data (Figure 2b – f). For example, on the two timsTOF Ultra 2 global proteome datasets, we observed that fine-tuning the models improved Spearman correlations for 54.53% and 61.90% of the peptides, respectively (Figure 2b – c). For example, the fragment ion intensity prediction for the peptide IAEPEPEPVPTLQAK detected in the first global proteome dataset before and after fine-tuning is shown in Figure 2e. Notably, a much larger improvement was observed on the phosphoproteome DIA dataset, where 79.92% of the peptides showed improved Spearman correlations after fine-tuning (Figure 2d). An example of predictions for the phosphopeptide IGPLGLsPK, in which the lowercase s indicates the phosphorylated serine residue, is shown in Figure 2f.

Next, we evaluated the performance of RT fine-tuning by repeating a similar setup (train on human and test on yeast) on the same three timsTOF DIA datasets generated using two different LC settings. It is worth noting that, unlike peptide fragment ion intensity, the retention time of a peptide is determined by LC systems rather than MS instruments. Therefore, from a methodological standpoint, RT model fine-tuning is MS instrument-agnostic and follows the same procedure for timsTOF and non-timsTOF DIA data. As shown in Figure 3a–c, RT models fine-tuned on DIA data consistently outperformed the pretrained DDA model, yielding higher squared correlation coefficients (*R*^2^). Consistent with our previous study [14], fine-tuned RT models showed stronger linear agreement and tighter RT accuracy across datasets, whereas the pretrained DDA model exhibited nonlinearity in the extremes of the RT range for all three datasets. Notably, fine-tuning enabled the models to effectively learn elution behavior for peptides eluting at the very beginning or end of the MS runs.

**Figure 3:**
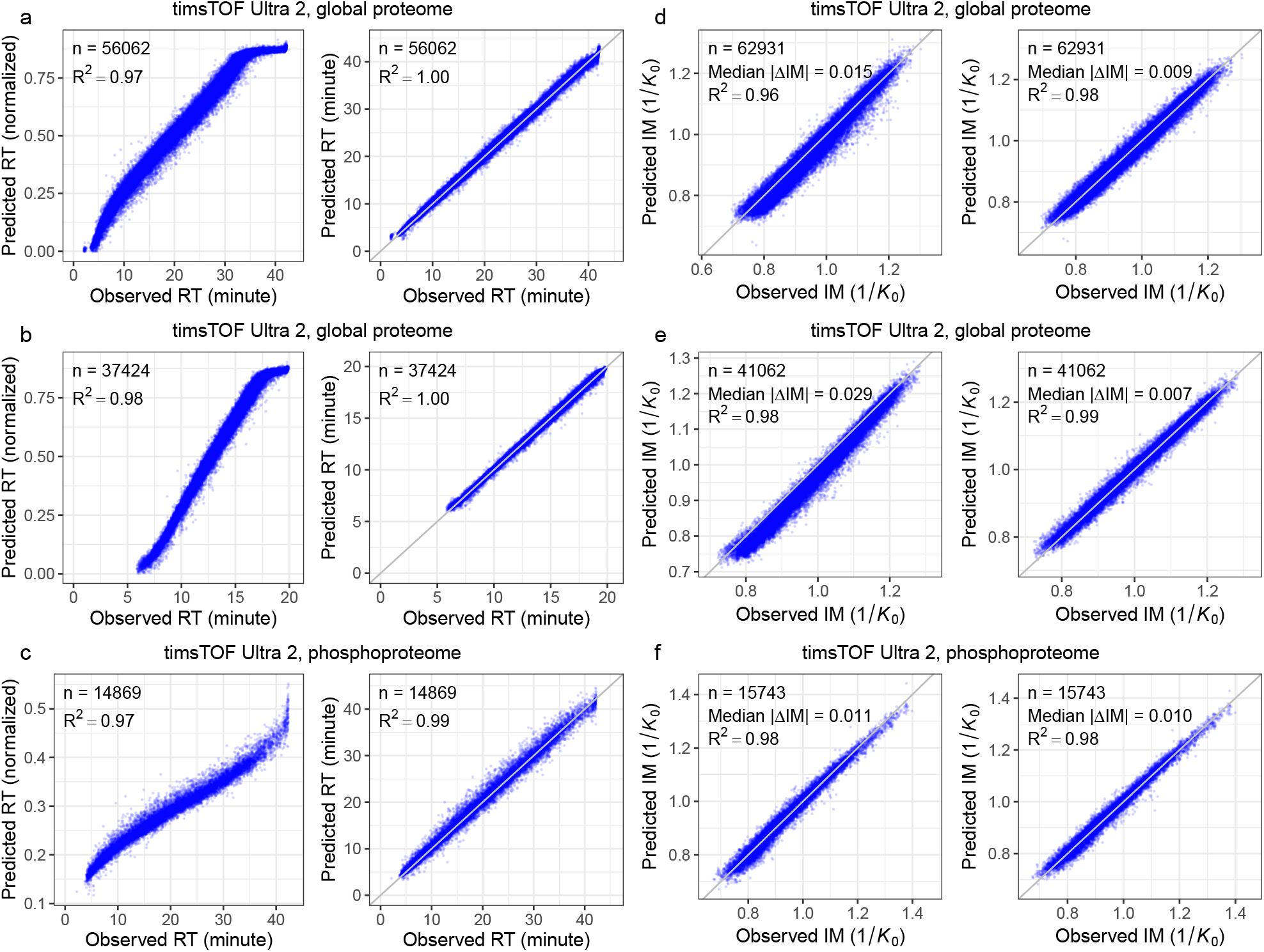
Retention time and ion mobility models fine-tuning evaluation. **(a–c)** Each panel plots observed RT versus RT predicted using the pretrained RT model (left panel) or fine-tuned RT model (right panel) in Carafe2. On each dataset, the fine-tuned RT model was trained using the human DIA data. Both types of RT models were evaluated using the corresponding yeast DIA data. The pretrained model predicts RT on a normalized scale between 0 and 1, whereas the fine-tuned models predict RT on an absolute scale aligned with the experimental RTs from the training DIA data. **(d–f)** Similar to **(a–c)**, each panel plots observed ion mobility versus predicted ion mobility (1*/K*_0_) using the pretrained model (left) or the fine-tuned model (right). In each plot, n is the number of peptides (RT) or precursors (ion mobility) used in evaluation. The squared correlation coefficient (*R*^2^) is shown in each panel. For ion mobility evaluation, the median absolute error is also shown.

Finally, we evaluated the fine-tuning of the ion mobility prediction model. Ion mobility values can vary systematically across timsTOF DIA datasets due to differences in MS instrument configuration and acquisition conditions, which could lead to experiment-specific shifts that are not captured by a single pretrained IM model. To quantify this effect, we compared ion mobility values for peptide precursors matched between two runs. This cross-study comparison reveals substantially larger dispersion around the *y* = *x* line than the within-study replicate comparison (Supplementary Figure 2a–b), indicating reduced agreement when transferring ion mobility annotations between studies. This cross-study variability motivates fine-tuning on experiment-specific DIA data: calibrating the prediction model to the target dataset is expected to better match the observed mobility scale and thereby improve the accuracy of the ion mobility annotations used for spectral library generation and ultimately improve peptide detection. As shown in Figure 3d–f and Supplementary Figure 3a –c, across all the three timsTOF DIA datasets, the Carafe2 fine-tuned IM models consistently outperformed the pretrained DDA model, demonstrating the benefit of DIA-specific fine-tuning for IM prediction on timsTOF data. Although fine-tuning improves ion mobility prediction relative to the pretrained DDA model, the residual prediction error (median absolute prediction error: 0.007–0.010) remains obviously larger than the ion mobility deviations (median absolute prediction deviation: 0.002) observed between replicate runs (Supplementary Figure 2–3), indicating that there is still room for further improvement. In addition, the cross-run variability analysis suggests that increasing training data size by combining peptides detected from multiple studies using different timsTOF instruments to improve ion mobility prediction will be challenging due to the ion mobility drifts across studies.

### 2.3 Carafe2 increases the number of peptides detected with DIA-NN

We next assessed how DIA fine-tuning in Carafe2 — applied to predicted retention time (RT), fragment ion intensities, and ion mobility — contributes to statistical power for peptide detection in DIA-NN. In this analysis, as illustrated in Figure 4a, we used human timsTOF DIA data as the source of training data and tested the performance using yeast DIA data generated under the same LC/MS condition. Performance was then quantified as the number of peptide precursors detected subject to specific criteria (such as 1% precursor-level FDR for global proteome DIA data; see Methods section for details). As shown in Figure 4b and Supplementary Figure 4a, using a global proteome DIA dataset generated using a timsTOF Ultra 2 instrument, incorporating Carafe2-predicted retention time (noted as “RT” in the figure) alone increased detections relative to the DDA-trained baseline library by 8.02% (from 65,118 to 70,342 precursors). Similarly, using predicted fragment ion intensities (MS2) yielded an increase of 7.75% (to 70,166 precursors). Adding predicted ion mobility alone provided a smaller but consistent gain (67,022; +2.92%). Notably, fine-tuning all the three attributes (noted as “Carafe2” in the figure) delivered the largest gain, increasing detections to 73,389 precursors (+12.70%), suggesting that accuracy gains in RT, MS2, and IM predictions are complementary, with the largest improvement achieved when all three are fine-tuned. Comparing with the *in silico* spectral library generated by DIA-NN (i.e., performing the search using DIA-NN’s “library-free” mode), 5.39% more precursors were detected using the Carafe2 fully fine-tuned library. We repeated this analysis on two additional timsTOF DIA datasets: a global proteome dataset generated previously from a different laboratory using the same type of timsTOF instrument but with a shorter LC gradient setting and a phosphoproteome DIA dataset (Figure 4c –d and Supplementary Figure 4b –c). The same trend was observed on both datasets. Notably, fragment ion intensity fine-tuning improved peptide detection much more on the phosphoproteomics dataset than on the other datasets, consistent with its stronger gains in fragment ion intensity prediction (Figure 2d). These results show that fine-tuning improves peptide detection with DIA-NN and suggest that fine-tuning is particularly valuable when the pre-trained models have limited representation of the target experimental conditions, enabling substantial gains by adapting predictions to the specific instrument and acquisition setting.

**Figure 4:**
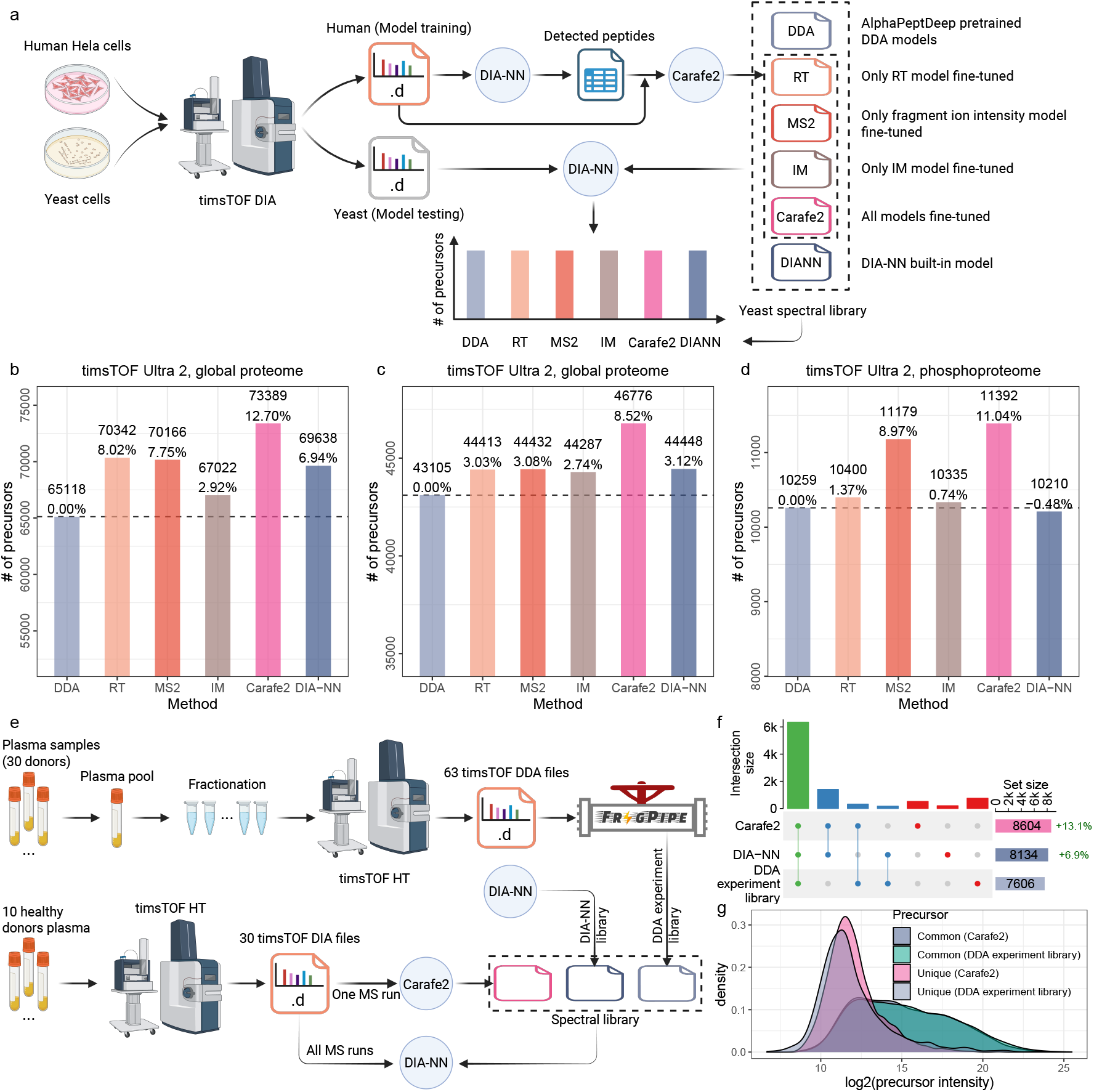
Comparing in silico spectral libraries generated by different methods. **(a)** The overview of *in silico* library generation methods and evaluation. **(b)** Peptide precursors comparisons on a yeast dataset generated on a timsTOF Ultra 2 instrument. **(c)** Peptide precursors comparisons on a yeast dataset generated on a different timsTOF Ultra 2 instrument with shorter LC gradient. **(d)** Phosphopeptide precursors comparisons on a yeast phosphoproteome DIA dataset generated on a timsTOF Ultra 2 instrument. Each dataset includes human DIA data as training data which was generated using the same LC and MS settings as the test yeast data. For the bar plots, for each dataset, the number of precursors accepted at the 1% precursor-level FDR are shown. The number on each bar is the percent improvement when comparing to the library generated using the AlphaPeptDeep pretrained DDA models. For x-axis, DDA: spectral library generated using the pretrained DDA model from AlphaPeptDeep; MS2: spectral library generated using a fine-tuned fragment ion intensity prediction model but the pretrained RT and IM models from AlphaPept-Deep; RT: spectral library generated using a fine-tuned RT prediction model but the pretrained fragment ion intensity and IM models from AlphaPeptDeep; IM: spectral library generated using a fine-tuned IM prediction model but the pretrained fragment ion intensity and RT models from AlphaPeptDeep; Carafe2: spectral library generated using fine-tuned fragment ion intensity, RT and IM prediction models; DIA-NN: spectral library generated using DIA-NN’s built-in model. **(e)** Comparing experiment spectral library with *in silico* spectral libraries on a human plasma timsTOF DIA dataset. The DDA experiment spectral library was generated using FragPipe on 63 DDA timsTOF runs from the same study. The *in silico* spectral libraries were generated using DIA-NN and Carafe2, respectively. **(f)** An upset plot showing the number of precursors detected using different spectral libraries. **(g)** A precu9rsor intensity density plot showing the distribution of precursor intensities for precursors detected between Carafe2 and DDA experiment library. Panels **(a)** and **(e)** were created using BioRender.

To further demonstrate the utility of the experiment-specific *in silico* spectral libraries generated by Carafe2, we reanalyzed a recently published DIA dataset of human plasma samples generated using a timsTOF HT instrument, comprising 30 DIA-MS runs [26]. In this analysis, as illustrated in Figure 4e, three spectral libraries were compared: (1) a Carafe2-generated library fine-tuned on a plasma DIA MS run from the same study, (2) an *in silico* spectral library generated using DIA-NN, and (3) an experimental library generated using FragPipe on 63 DDA timsTOF runs generated from the same study. As shown in Figure 4f, the Carafe2-generated library enabled the detection of more precursors compared to both the experimental spectral library derived from DDA data and the DIA-NN generated *in silico* spectral library. Specifically, Carafe2 identified 13.1% more peptide precursors than the experimental library and 5.8% more peptide precursors than the DIA-NN library at 1% precursor-level FDR. Although the precursors uniquely detected using Carafe2 had lower intensities than those detected by both Carafe2 and the DDA experimental library (Figure 4g), they tended to have higher intensities than the precursors uniquely detected by the DDA experimental library, suggesting that the Carafe2-specific detections are not of lower quality based on intensity distribution alone. Importantly, both *in silico* libraries exceeded the performance of the DDA-derived experimental library constructed from 63 DDA runs, highlighting the advantage of *in silico* libraries for complex plasma samples, with Carafe2 achieving the best overall performance. We repeated this analysis using a different timsTOF dataset generated from human triple-negative breast cancer samples [27], and observed the same trend: the Carafe2 fine-tuned library detected 45.5% more precursors than the DDA experimental library and 7.5% more than the DIA-NN generated *in silico* library (Supplementary Figure 5).

As an additional assessment of the quality of the *in silico* spectral libraries generated by Carafe2, we tested whether DIA fine-tuning with Carafe2 introduces any bias that could inflate FDR estimates specifically at the precursor level. To this end, we evaluated precursor-level FDR control on a global proteome timsTOF DIA dataset using an entrapment strategy (Figure 5a), employing two different techniques that yield an upper bound and a lower bound, as described in our previous studies [14, 28]. We observed that the FDR control is similar between using spectral libraries generated with the DDA data pretrained models (Figure 5b–c) and fine-tuned spectral libraries generated by Carafe2 (Figure 5d–f). In particular, when we analyzed the same human DIA data and compared the estimated false discovery proportion (FDP) using the upper bound method for DIA-NN with three different spectral libraries at the 1% FDR threshold reported by DIA-NN, we obtained estimates of 0.83% using the spectral library generated by the pretrained DDA models (Figure 5b), 0.97% using the spectral library generated by DIA-NN (Figure 5c), and 0.95% using a Carafe2 fine-tuned spectral library trained on a yeast DIA run from the same experiment (Figure 5d). Furthermore, the estimated FDP was 0.97% using a Carafe2 fine-tuned spectral library trained on a replicate DIA run (Figure 5e), while the estimated FDP was also 0.97% using a Carafe2 fine-tuned spectral library trained on the same DIA run (Figure 5f). Importantly, all the estimated upper bound of FDPs were under 1%. These results mirror our previous analysis on a TripleTOF 5600 DIA dataset [14] and indicate that using Carafe2 fine-tuned spectral libraries does not compromise precursor-level FDR control in DIA-NN.

**Figure 5:**
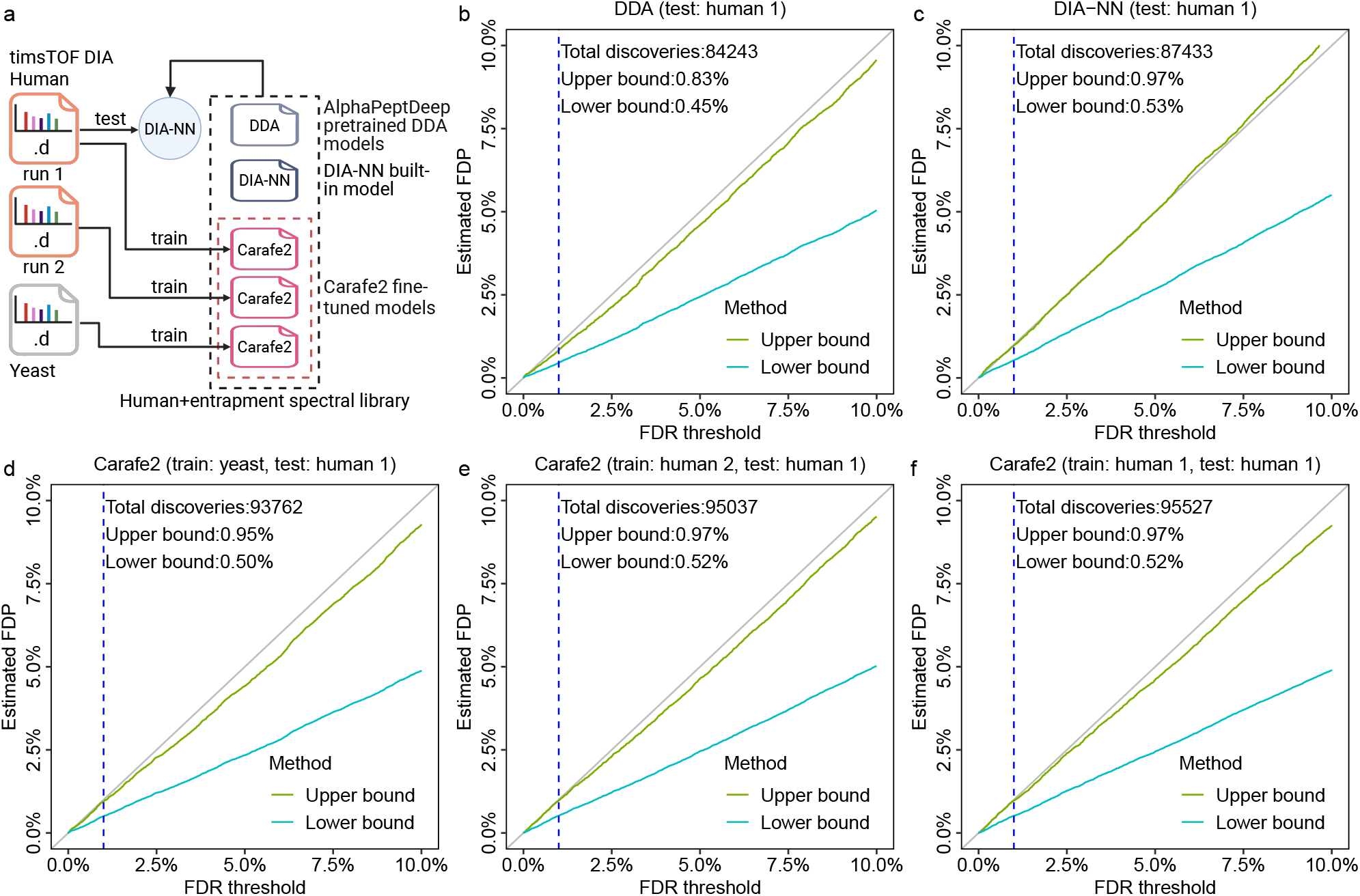
FDR control evaluation. **(a)** FDR control evaluation was performed using an entrapment strategy on a timsTOF Ultra 2 human DIA dataset using five different *in silico* spectral libraries: **(b)** a spectral library generated using the pretrained DDA models from AlphaPeptDeep; **(c)** a spectral library generated using DIA-NN’s built-in model; **(d)** a spectral library generated using Carafe2 with models fine-tuned on a yeast DIA run from the same experiment; **(e)** a spectral library generated using Carafe2 with models fine-tuned on a replicate human DIA run from the same experiment, and **(f)** a spectral library generated using Carafe2 with models fine-tuned on the same human DIA run. The x-axis shows different FDR thresholds reported by DIA-NN while the y-axis shows the estimated FDPs using the upper bound and lower bound methods. The dashed vertical line is at the 1% FDR threshold, as are the numbers reported in the text in the figure. Panel **(a)** was created using BioRender.

### 2.4 Quantification evaluation

To benchmark quantification performance using Carafe2, we analyzed a mixed-species dataset (PXD062685) acquired on a timsTOF SCP instrument and compared Carafe2’s fully fine-tuned library against a few alternative spectral libraries (Figure 6a). Each run in the dataset is derived from one of the two samples, A and B, each containing a mixture of human, yeast, and *E*.*coli* peptides at different ratios (1:1 for human, 2:1 for yeast and 1:4 for *E*.*coli*) with three technical replicates each. As shown in Figure 6b, using the Carafe2 fully fine-tuned library increased peptide detections, yielding 8.79% more precursors than the library generated using the AlphaPeptDeep DDA-pretrained models and 9.03% more than the library generated using DIA-NN built-in model (Figure 6b). This observation further supports the conclusion that jointly refining RT, fragment ion intensity, and IM predictions improves precursor detection in DIA-NN.

**Figure 6:**
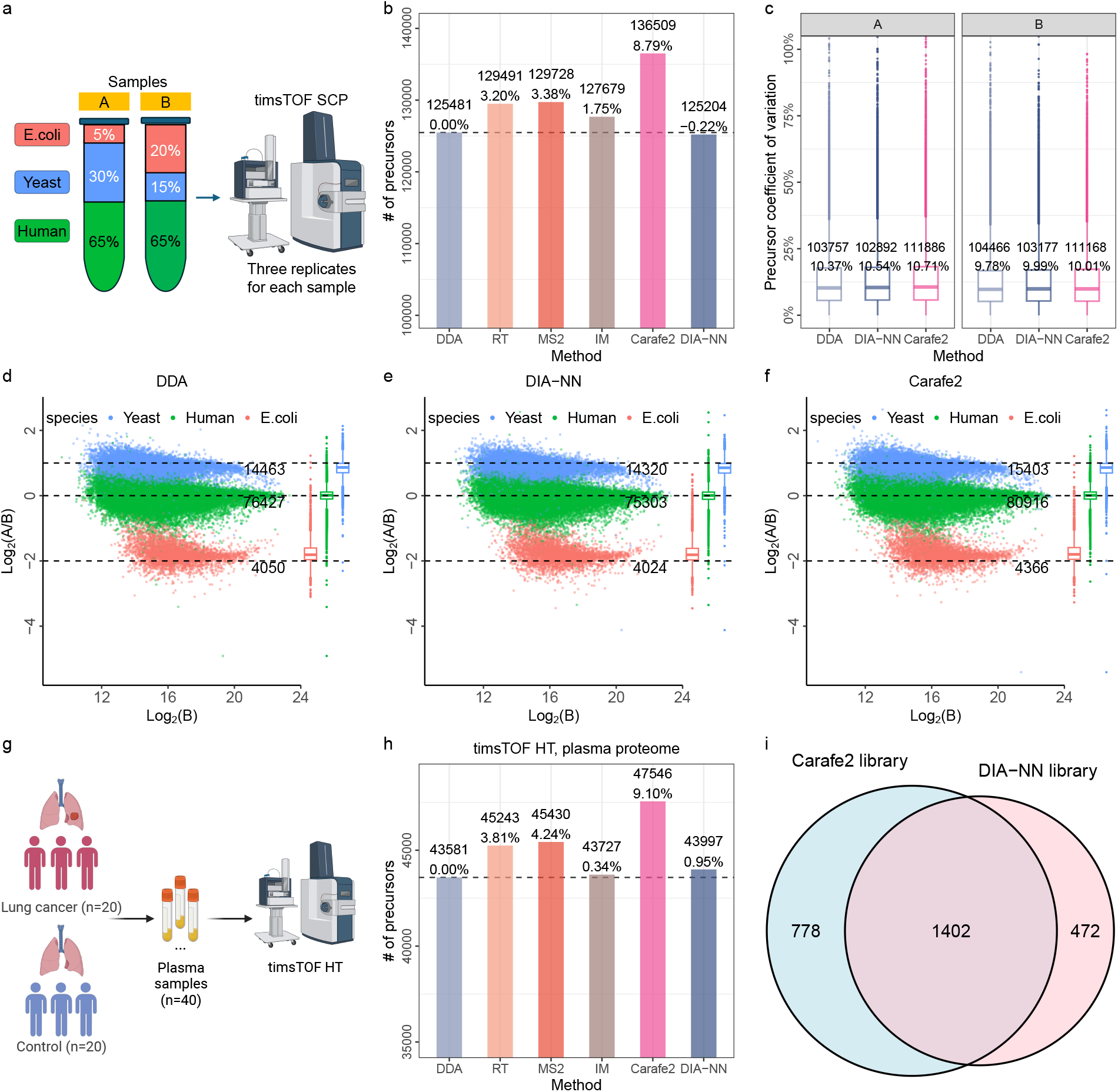
Quantification evaluation of Carafe2 with DIA-NN. Quantitative evaluation using a mixed-species dataset. **(a)** An overview of the dataset (two samples: A and B, three technical replicates each). **(b)** Comparing peptide precursors detected using six different *in silico* spectral libraries. **(c)** The distribution of CVs of precursor intensity for samples A and B. Numbers in the first row of the boxplots are the numbers of peptide precursors quantified. Numbers in the second row are the median CVs. **(d–f)** Log-transformed ratios (log2(A/B)) of peptide precursors over log-transformed intensities of sample B for the AlphaPeptDeep DDA-pretrained models **(d)**, the DIA-NN built-in model (**(e)**) and the Carafe2 fully fine-tuned models (**(f)**) generated spectral libraries. Dashed black lines indicate the expected log2(A/B) ratios: 1:1 for human, 2:1 for yeast and 1:4 for *E*.*coli*. The number under each line is the number of precursors quantified from a species used in the plot. The boxplot on the right side of each plot shows the distribution of the log2(A/B) ratios. **(g)** An overview of the lung cancer plasma dataset. **(h)** Comparing peptide precursors detected using six different *in silico* spectral libraries on the lung cancer plasma dataset. **(i)** Comparing differentially expressed precursors detected using Carafe fully fine-tuned and DIA-NN libraries. The p-values were calculated using a two-tailed t-test on the log2 transformed quantification values with Benjamini-Hochberg correction. Part of panel **(a)** and the full panel **(g)** were created using BioRender.

To evaluate quantification precision, we examined replicate consistency of peptide precursor intensities for samples A and B, computing the coefficient of variation (CV) of precursors across three technical replicates and comparing Carafe2 to the AlphaPeptDeep DDA-pretrained and DIA-NN libraries. As shown in Figure 6c, Carafe2 fully fine-tuned library produced median CVs similar to the other libraries, indicating comparable precursor-level precision.

Next, we evaluated quantification accuracy by comparing measured precursor ratios between samples A and B to the expected mixture ratios (1:1 human, 2:1 yeast, 1:4 *E. coli*). Across species, Carafe2 produced ratio estimates that closely tracked expectations and were comparable to those from the AlphaPeptDeep DDA-pretrained models and DIA-NN built-in model generated libraries (Figures 6d–f).

To further assess the quantification performance of Carafe2 in a biological context, we evaluated its ability to detect differentially expressed precursors using a clinical human lung cancer plasma dataset (Figure 6g). The dataset comprises 40 DIA runs corresponding to 20 lung cancer and 20 control samples. We first compared the number of peptide precursors detected using six different spectral libraries. Consistent with our previous results, the fully fine-tuned Carafe2 library outperformed all other libraries, yielding the highest number of precursor detections (Figure 6h). We next performed differential expression analysis to compare the Carafe2 fine-tuned library against the DIA-NN built-in library. As shown in Figure 6i, the Carafe2 library identified 16.3% more significantly regulated precursors between the cancer and control groups compared to the DIA-NN library. These results demonstrate that the improved peptide detection and quantification precision provided by Carafe2 translates directly into increased statistical power for discovering differentially expressed biological features.

### 2.5 Timsviewer enables visualization of timsTOF DIA data with fine-tuned spectral libraries from Carafe2

To facilitate the inspection of timsTOF DIA data analyzed with *in silico* spectral libraries generated by Carafe2, we developed Timsviewer, a standalone visualization tool written in Rust. Designed for broad accessibility, Timsviewer is cross-platform and supports Linux, Windows, and macOS. As shown in Figure 7, Timsviewer allows users to visualize the extracted ion chromatograms (XICs) for both peptide precursors and their corresponding fragment ions from Carafe2-generated spectral libraries. Notably, the tool can directly read timsTOF raw data without relying on third-party software developer kits. Upon loading a library, Timsviewer automatically indexes the raw data — a process that typically completes in less than three minutes — enabling real-time extraction and visualization of any precursor within the library. Timsviewer also includes an MS2 spectrum panel, which shows a mirror plot comparing observed fragment ion intensities at a specific time point with predicted fragment ion intensities from the loaded spectral library. The tool thus provides an intuitive interface for exploring the data, enabling users to assess the quality of peptide detections and the performance of the spectral library. Timsviewer supports various features, including zooming and panning, making it a valuable resource for researchers working with timsTOF DIA data.

**Figure 7:**
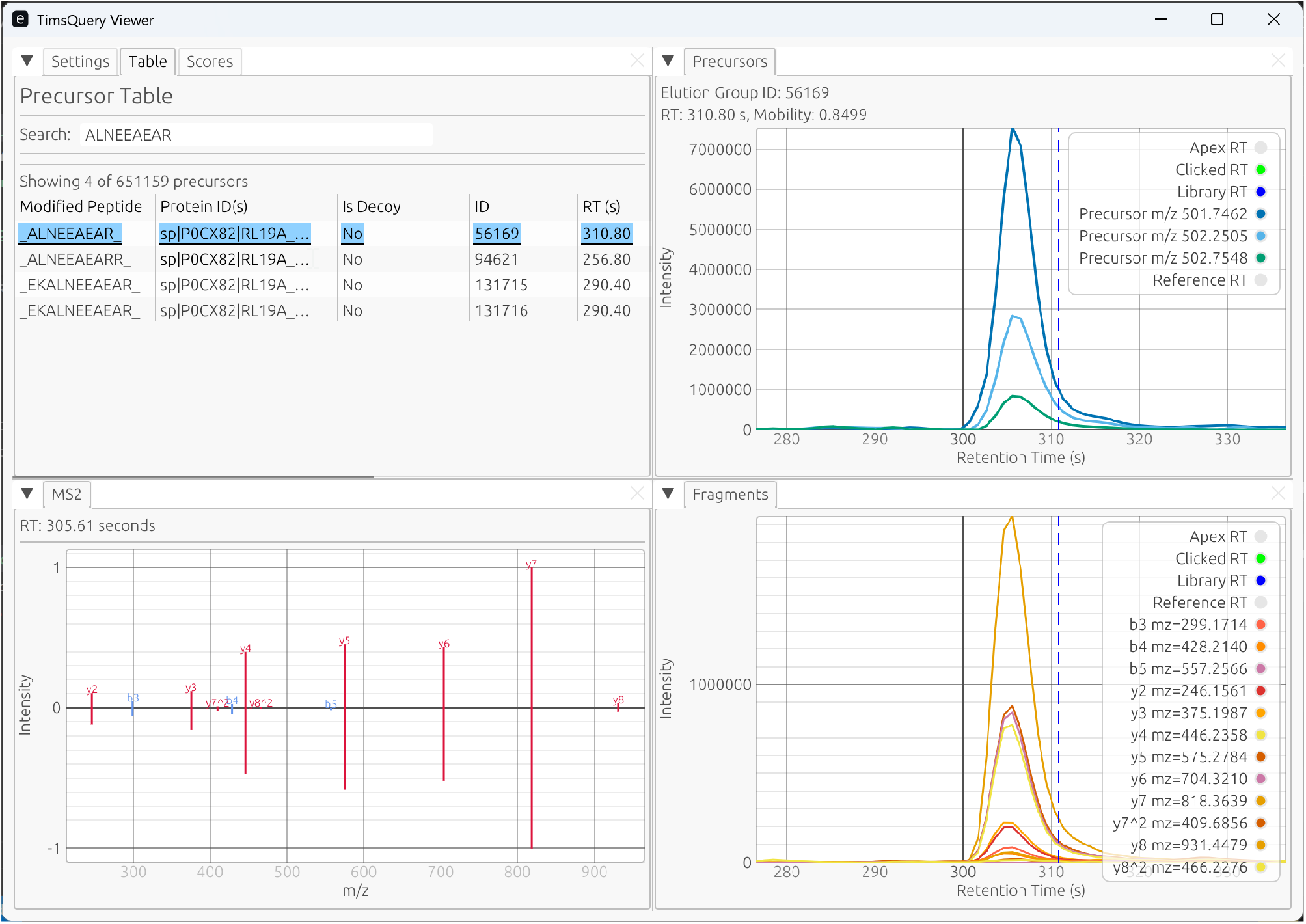
Timsviewer interface. The Timsviewer interface consists of five main panels: parameter setting panel, precursor table panel, precursor XIC panel, fragment ion XIC panel, and MS2 spectrum panel. The precursor table panel shows a list of precursors in the spectral library, with each row representing a unique precursor. The precursor and fragment ion panels show the extracted ion chromatograms for each precursor and its corresponding fragment ions, respectively. The MS2 spectrum panel shows the observed (top panel) and predicted (bottom panel) fragment ion intensities for a selected precursor at a specific time point.

## 3 Discussion

In this study, we introduced Carafe2, an extension of Carafe that enables generation of experiment-specific *in silico* spectral libraries for timsTOF DIA proteomics. A key design goal of Carafe2 is to reduce preprocessing overhead while fully exploiting the additional ion mobility dimension. To that end, Carafe2 operates on native timsTOF DIA raw data (Bruker .d directories) via TimsQuery, enabling efficient access to MS signals without conversion to intermediate MS formats. Across diverse timsTOF DIA datasets, we showed that DIA-based fine-tuning improves the accuracy of retention time, fragment ion intensity, and ion mobility predictions, and that these improvements translate into increased statistical power for peptide detection when used with DIA-NN. To support practical adoption, we provide a graphical user interface that streamlines common workflows, including library generation from existing DIA-NN results, library generation with DIA-NN search executed within the same workflow, and end-to-end DIA analysis using Carafe2 and DIA-NN. Because Carafe2 is integrated into Skyline and provides experiment-specific predictions of retention time, fragment ion intensity, and ion mobility for each precursor, its utility extends beyond DIA library generation to targeted proteomics applications. In particular, these highly tailored predictions can guide targeted method design and support data analysis in Skyline by informing optimal peptide and transition selection, accurate retention time scheduling, and ion mobility-aware assay refinement. To support the interpretation of results — such as manually verifying the quality of key peptide detections used for downstream biological validation — we additionally developed Timsviewer, a standalone tool for interactive inspection of timsTOF DIA data alongside Carafe2-generated spectral libraries. This tool is particularly important because accurate FDR control is still challenging for DIA data analysis [28, 29]. Together, these components provide an end-to-end, open-source toolkit for building, applying, and evaluating high-quality spectral libraries tailored to specific timsTOF DIA experiments.

Despite these advances, several limitations point to clear opportunities for future work. First, although fine-tuning improves prediction accuracy, residual errors in IM predictions indicate that model performance could be further improved, for example through additional training data, improved training strategies, or more advanced model architectures. Second, library generation can be computationally demanding, particularly for analyses with extremely large search spaces (e.g., metaproteomics databases or non-specific searches for immunopeptidomics). Accordingly, further engineering and algorithmic optimizations will be important to improve scalability and throughput. Third, future work could incorporate additional predictive modules, such as precursor charge state prediction and peptide detectability prediction, to prioritize likely observable precursors and thereby optimize (or reduce) library size. Finally, although we demonstrated benefits using DIA-NN, it will be valuable to assess the generality of Carafe2-generated libraries with additional DIA analysis tools. Additionally, Timsviewer could be extended with support for additional MS data formats (e.g., mzML and Thermo raw files), further streamlining exploratory data analysis and quality control.

## 4 Methods

### 4.1 Deep learning models

The deep learning model architectures used in Carafe2 for predicting RT and fragment ion intensity are based on the models previously described in Carafe [14]. For ion mobility (IM) prediction, we adopted the architecture from AlphaPeptDeep [13]. The AlphaPeptDeep IM model is also referred to as a CCS model since the model directly predicts CCS, but there is an internal conversion between ion mobility (1*/K*_0_) and CCS. The model consists of an embedding layer for sequence, modifications and charge states, a convolutional neural network (CNN) layer, followed by two bidirectional long short-term memory (BiLSTM) layers. The training parameters were set as follows: epoch, 40; warmup epoch, 10; learning rate, 0.0001; batch size, 1024. L1 loss was used for IM model fine-tuning. The training parameters for RT and fragment ion intensity models were the same as those used in the previous Carafe [14].

### 4.2 Training and testing data generation

We trained three different types of prediction models (fragment ion intensity, RT, and ion mobility) for each dataset. Following our previous work [14], unless otherwise specified we trained the models using a DIA run from one species and then tested the models on DIA data from a different species. For each training or testing DIA run, we used DIA-NN (version 2.3.2) in library-free mode to detect peptides at 1% precursor-level FDR.

To generate training data for fragment ion intensity prediction model training, we followed the general strategies described in Carafe [14] with the following updates for supporting timsTOF DIA data. First, for each detected precursor accepted at 1% FDR, we generated its theoretical fragment ions (e.g., b/y ions with charge states up to 2) and then used TimsQuery (https://github.com/TalusBio/timsbuktoolkit/) to extract the corresponding ion chromatogram (XIC) from the training MS raw data based on the precursor’s detected apex RT and ion mobility, as determined by DIA-NN, with the following parameters: a specified fragment ion *m/z* tolerance (e.g., 20 ppm), a specified RT window and ion mobility tolerance (3% by default). Each query returned a json object in which the XIC data points were stored. Next, the apex retention time and peak boundaries for the precursor were refined using a procedure similar to that described previously [14]. Finally, for each precursor, we extracted its intensity at the refined apex retention time for each fragment ion as the observed fragment ion intensity for model training. For fragment ion intensity model training, the collision energy associated with each precursor was extracted from the corresponding training DIA run. To identify fragment ions that were interfered by co-eluting peptides, we used the previously described peptide-centric interference detection algorithm [14].

### 4.3 Data extraction from native timsTOF raw data using TimsQuery

The application initially preprocesses the raw data with a variant of the DBSCAN algorithm [30], modified to account for the intensity dimension in the data. Briefly, within each frame, low intensity peaks are added to the highest intensity peak within their immediate neighborhood (as defined by a tolerance setting in the mobility and *m/z* dimension, which default to 5% and 5 ppm respectively). This process is carried out iteratively until all pairs of neighboring peaks have been aggregated. The final mobility and *m/z* values are calculated from the weighted average of all the aggregated peaks for a given cluster. After clustering, the peaks are organized per window group in a data structure similar to a 2-level b+ tree, where the initial level is indexed by *m/z* and the second level is sorted by retention time. Similar data structures have been used in the past for mass spectrometry data, notably done by MSFragger [25] and Sage [31] for their search space. This data structure allows efficient querying of full chromatograms with four binary search operations on average (and less than a hundred boolean operations).

### 4.4 Proteome and phosphoproteome sample preparation

The samples for HeLa and yeast global proteome were prepared analogously to Wen *et al*. 2025 [14] using the protein aggregate capture method [32, 33]. Samples for the human phosphoproteome were prepared as described previously [14, 34–36]. Samples for the yeast phosphoproteome were prepared in the similar manner as Wen *et al*. 2025 [14] except that the yeast strain differed. Here we used a BY4741 yeast strain expressing a beta-estradiol inducible v-Src kinase gifted from the Landry lab [37].

### 4.5 timsTOF data generation

Peptides were analyzed with an Evosep One coupled with a Bruker timsTOF Ultra II mass spectrometer. We used an Evosep EV1106 column (15 cm *×* 150 *µ*m, packed with Dr Maisch ReproSil-Pur C18, 1.9 *µ*m beads). Solvent A was 0.1% formic acid in water and solvent B was 0.1% formic acid in acetonitrile. For each injection, 100 ng of peptides were loaded onto an Evotip and were separated using the Evosep 30 samples per day (SPD) method. Mass spectrometry diaPASEF measurements were acquired using 25 m/z isolation windows spanning a range from 400 to 1000 m/z and 8×3 TIMS ramps spaced from 0.64 to 1.45 1*/K*_0_. The accumulation and ramp times were 75ms and ion charge control was turned on. For phosphoproteomics, 2×12 TIMS ramps were used with variable isolation width, spanning 400 to 1400 m/z and 0.60 to 1.60 1*/K*_0_. The accumulation and ramp times were 75ms and ion charge control was also turned on.

### 4.6 Protein databases

Protein sequences for human (UP000005640, containing 20,597 proteins) and yeast (UP000002311, containing 6,060 proteins) were downloaded from UniProt (02/2024). These are the same protein databases used in our previous Carafe study [14]. The mixed species (human, yeast and *E*.*coli*) protein database was downloaded from PRIDE with accession number PXD062685.

### 4.7 *In silico* spectral library generation using Carafe2

To generate an experiment-specific *in silico* spectral library using Carafe2 for a specific instrument setting, one or more timsTOF DIA MS runs were used as training data and were analyzed using DIA-NN (version 2.3.2). Then peptides detected in the training data were used for model fine-tuning. In the study, for a given protein database, we first performed *in silico* protein digestion using trypsin without proline suppression with a maximum of one missed cleavage site allowed. Peptides with length between 7 and 35 amino acids were considered and precursor charge states from 2+ to 4+ were considered. The fixed modifications were set as Carbamidomethyl (C) by default. No variable modifications were applied for global proteome data while phosphorylation (STY) was set as variable modification for phosphoproteome data by default with a maximum of one variable modification allowed. Finally, we applied the trained models to predict fragment ion intensities, RTs and ion mobility values for all the peptidoforms to generate the spectral libraries. During fragment ion intensity prediction, the collision energy for each precursor was set to the value assigned to the isolation window containing that precursor in the training DIA run. During spectral library generation, only precursors and fragment ions with m/z values within the m/z scan range of the corresponding DIA data used for model training were included in each library. For each precursor, the top 20 fragment ions were selected based on the predicted fragment ion intensities.

### 4.8 DIA-NN analysis

DIA-NN (version 2.3.2) [23] analysis was performed using the following parameters: fixed modification, carbamidomethyl (C); no variable modification was set except phosphorylation was set for the phosphoproteomics data; enzyme, Trypsin/P with one missed cleavage site allowed; peptide length range, 7–35; precursor charge range, 2+ to 4+. The setting of “N-term M excision” was disabled; library generation, “IDs, RT & IM profiling”. For the phosphoproteomics data, the maximum number of variable modifications was set to 1. Peptidoform scoring was also enabled for the phosphoproteomics data analysis. Both parameters “Mass accuracy” and “MS1 accuracy” were set to 15 ppm. The precursor FDR threshold was set to 1%. For single-run DIA data, the “Q.Value” from the main report was used as precursor q-value for downstream analysis. For datasets with multiple runs, the “Lib.Q.Value” from the main report was used as precursor q-value for downstream analysis. For phosphoproteomics data, two additional filters were applied: peptidoform q-value ≤ 0.01, and PTM site confidence ≥ 0.75.

### 4.9 Reanalysis of human plasma DIA data

The raw data of the human plasma DIA dataset (30 MS runs) generated using a timsTOF HT system published recently [26] were downloaded from PRIDE [38] with accession number PXD058337. A total of 63 DDA runs generated from the same study were also downloaded for building an experimental spectral library using FragPipe (version 24.0) [25, 39]. Specifically, the DDA timsTOF data were searched against the human protein database (UP000005640) using FragPipe with the “DIA SpecLib Quant” workflow. To be consistent with the DIA data analysis using DIA-NN, the following parameters were updated for the FragPipe analysis: one missed cleavage site allowed; peptide length range, 7–35; no variable modification was set. The spectral library file “library.tsv” generated by FragPipe was used for the DIA-NN analysis of the plasma DIA data. For the Carafe2-generated *in silico* spectral library, we fine-tuned the RT, fragment ion intensity, and IM prediction models using one DIA run (file name: “HC1-30SPD Repl1 S4-A1 1 8993.d”) from the plasma DIA data. The Carafe2-generated library and the DIA-NN built-in *in silico* library were also used for the DIA-NN analysis of the plasma DIA data. The settings for the DIA-NN analysis were the same as described above.

### 4.10 Reanalysis of triple-negative breast cancer DIA data

The raw data of the human triple-negative breast cancer (TNBC) DIA dataset (16 MS runs) generated using a timsTOF Pro system published previously [27] were downloaded from PRIDE [38] with accession number PXD047793. A total of 12 DDA runs generated from a pool of 105 TNBC tissues from the same study were also downloaded for building an experimental spectral library using FragPipe (version 24.0) [25, 39]. Specifically, the DDA timsTOF data were searched against the human protein database (UP000005640) using FragPipe with the “DIA SpecLib Quant” workflow. To be consistent with the DIA data analysis using DIA-NN, the following parameters were updated for the FragPipe analysis: one missed cleavage site allowed; peptide length range, 7–35; no variable modification was set. The spectral library file “library.tsv” generated by FragPipe was used for the DIA-NN analysis of the DIA data. For the Carafe2-generated *in silico* spectral library, we fine-tuned the RT, fragment ion intensity, and IM prediction models using one DIA run (file name: “4223_TIMS2_001_4_Slot2-1_1_2869.d”) from the DIA data. The Carafe2-generated library and the DIA-NN built-in *in silico* library were also used for the DIA-NN analysis of the DIA data. The settings for the DIA-NN analysis were the same as described in the DIA-NN analysis section.

### 4.11 FDR control evaluation

FDRBench (version 0.0.1, https://github.com/Noble-Lab/FDRBench) [28] was used to generate a peptide-level entrapment database and estimate FDP for FDR control evaluation. In the evaluation, protein sequences for human (UP000005640, containing 20,597 proteins) were taken as the original target protein database. The original target proteins were *in silico* digested into peptides using trypsin (without proline suppression) with one missed cleavage allowed. Only peptides with lengths between 7 and 35 amino acids were considered. All “I” amino acids were converted to “L.” For each original target peptide, an entrapment peptide was generated using FDRBench with the C-terminal amino acid fixed. The final peptide level entrapment database was generated by combining the original target peptides and the entrapment peptides. FDP was estimated using the paired method (upper bound) and the lower bound method using FDRBench:

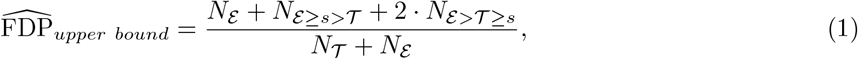

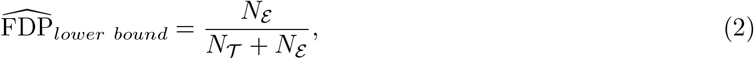

where *s* is the discovery cutoff score; *N*_ℰ≥*s>*𝒯_ denotes the number of discovered entrapment peptides (scoring ≥ *s*) for which their paired original target peptides score *< s*; and *N*_ℰ*>*𝒯≥ *s*_ is the number of discovered entrapment peptides for which the paired original target peptides scored lower but were still also discovered. *N*_𝒯_ and *N*_ℰ_ denote the number of original target and entrapment discoveries, respectively.

For the evaluation, five *in silico* spectral libraries were generated as shown in Figure 5a. One of them was generated using the pretrained DDA models from AlphaPeptDeep, one of them was generated using DIA-NN’s built-in model, and three of them were generated using Carafe2 with models fine-tuned on a yeast DIA run, a replicate human DIA run, and the same human DIA run that was used for testing, respectively. The parameters for library generation were similar as described in the previous section. The MS/MS data were searched against each spectral library using DIA-NN (version 2.3.2) with the following parameters: fixed modification, carbamidomethyl (C); no variable modification was set; enzyme digestion was disabled; peptide length range, 7–35; precursor charge range, 2+ to 4+; N terminal methionine excision was disabled. The precursor FDR was set to 10%. All other parameters were set to their default values. The column “Q.Value” from the main report generated by DIA-NN was used as precursor q-value for FDR control evaluation.

### 4.12 Quantification precision and accuracy evaluation

The mixed-species dataset was downloaded from PRIDE with accession number PXD062685. The MS/MS data were generated using a timsTOF SCP instrument. A total of six MS runs were included as shown in Figure 6a: three replicates for sample A and three replicates for sample B. In the dataset, peptides from three species (human, yeast and *E*.*coli*) were mixed together to generate samples A and B with expected A:B ratios of 1:1 for human, 2:1 for yeast and 1:4 for *E*.*coli* peptides. The first MS run from sample A from the dataset was used as the training MS run for Carafe2. Six spectral libraries were generated. Five of them were generated using Carafe2, and one of them was generated using DIA-NN’s built-in model. The parameters for library generation were the same as described in the library generation section. The MS/MS data were searched against each spectral library using DIA-NN (version 2.3.2) with the following parameters: fixed modification, carbamidomethyl (C); no variable modification was set; enzyme, Trypsin/P with one missed cleavage site allowed; peptide length range, 7–35; precursor charge range, 2+ to 4+; N terminal methionine excision was enabled. The column “Precursor.Normalised” from the main report generated by DIA-NN was used as precursor intensity for the quantification precision and accuracy evaluation.

### 4.13 Reanalysis of lung cancer plasma DIA data

The raw data of the human lung cancer plasma DIA dataset generated using a timsTOF HT system published previously [40] were downloaded from PRIDE [38] with accession number PXD047839. A total of 40 DIA runs generated from 20 lung cancer plasma samples and 20 control samples were used in the study. The plasma samples were processed using Seer’s Proteograph Assay Kit with a specific nanoparticle (noted as NP2 in the original study). For *in silico* spectral library generation using Carafe2, we fine-tuned the RT, fragment ion intensity, and IM prediction models using one DIA run (sample name: “HT-subject010-NP2”) from the DIA data. The Carafe2-generated libraries and the DIA-NN built-in *in silico* library were used for the DIA-NN analysis of the 40 DIA runs. The settings for the DIA-NN analysis were the same as described in the DIA-NN analysis section. For differential expression analysis, both the precursor level and protein group level quantification matrices generated by DIA-NN were used. Only precursors and proteins detected in at least 50% of samples in each cohort were included in the differential expression analysis. The fold changes comparing the cancer with the control group were log2 transformed. The p-values were calculated using a two-tailed t-test on the log2 transformed quantification values and were adjusted using the Benjamini-Hochberg procedure. Precursors and proteins with adjusted p-values ≤ 0.05 and log2 transformed fold changes ≥ *log*2(1.5) or ≤ −*log*2(1.5) were considered as significantly regulated.

### 4.14 Data availability

The MS/MS datasets generated in this study have been deposited to Panorama Public (ProteomeXchange identifier: PXD075483) and are available at https://panoramaweb.org/Carafe2.url. The mixed-species dataset was downloaded from PRIDE with accession number PXD062685. The timsTOF Ultra 2 with shorter LC gradient dataset was downloaded from PRIDE with accession number PXD070049. The human plasma dataset was downloaded from PRIDE with accession number PXD058337. The human triple-negative breast cancer dataset was downloaded from PRIDE with accession number PXD047793. The human lung cancer plasma dataset was downloaded from PRIDE with accession number PXD047839.

### 4.15 Code availability

Carafe2 is open source and is available with an Apache 2.0 license at https://github.com/Noble-Lab/Carafe. The Skyline version with Carafe2 integrated is available at https://skyline.ms/carafe.url. A Nextflow workflow for Carafe2 is available at https://nf-carafe-ai-ms.readthedocs.io. TimsQuery and Timsviewer are open source and are available with an Apache 2.0 license at https://github.com/Tal_usBio/timsbuktoolkit. Both Carafe2 and Timsviewer can be run on Linux, Windows, and macOS.

## Acknowledgments

This work was supported by National Science Foundation award 2245300, National Institutes of Health awards R24 GM141156, P30 AG013280, R35 GM152061, RM1 HG010461, the National Science Foundation Graduate Research Fellowship Program (Grant No. DGE-2140004, B.W.), and by the Intelligence Advanced Research Projects Activity (IARPA) TEI-REX program through the Army Research Office contract W911NF2220059. We gratefully acknowledge Lauren Fields for testing Carafe2 and help on figure generation.

## Author contributions

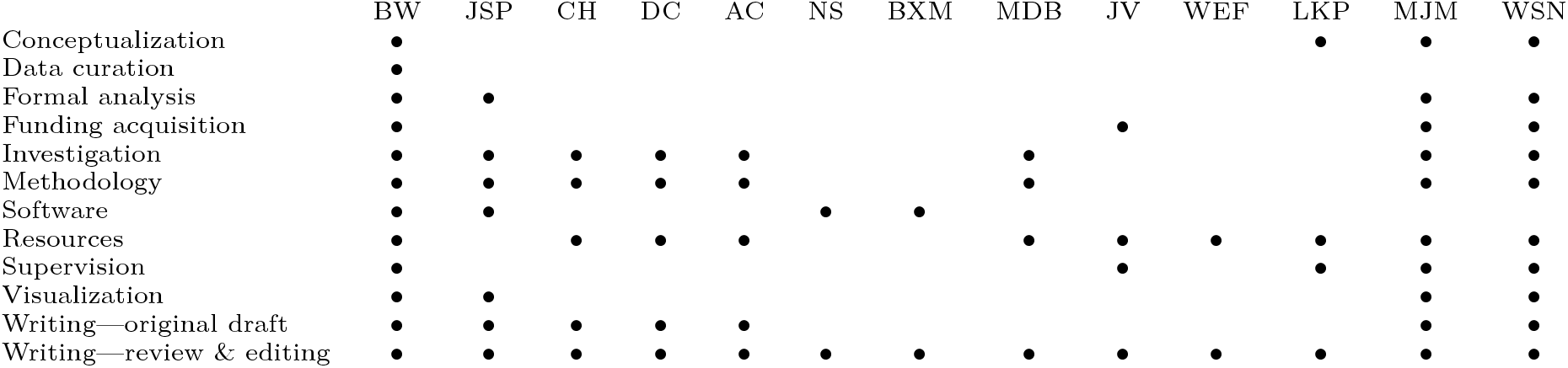

## Conflict of interest

The MacCoss Lab at the University of Washington receives funding from Agilent, Bruker, Sciex, Shimadzu, Thermo Fisher Scientific, and Waters to support the development of Skyline, a quantitative analysis software tool. MJM is a paid consultant for Thermo Fisher Scientific. WEF, JSP, DC, and LKP are employees and stakeholders of Talus Bioscience, a drug-discovery biotechnology company that develops, but does not currently sell software.

## Notes

https://github.com/Noble-Lab/Carafe

https://panoramaweb.org/Carafe2.url

