## Supplementary Figure for "Carafe2 enables high quality *in silico* spectral library generation for timsTOF data-independent acquisition proteomics"

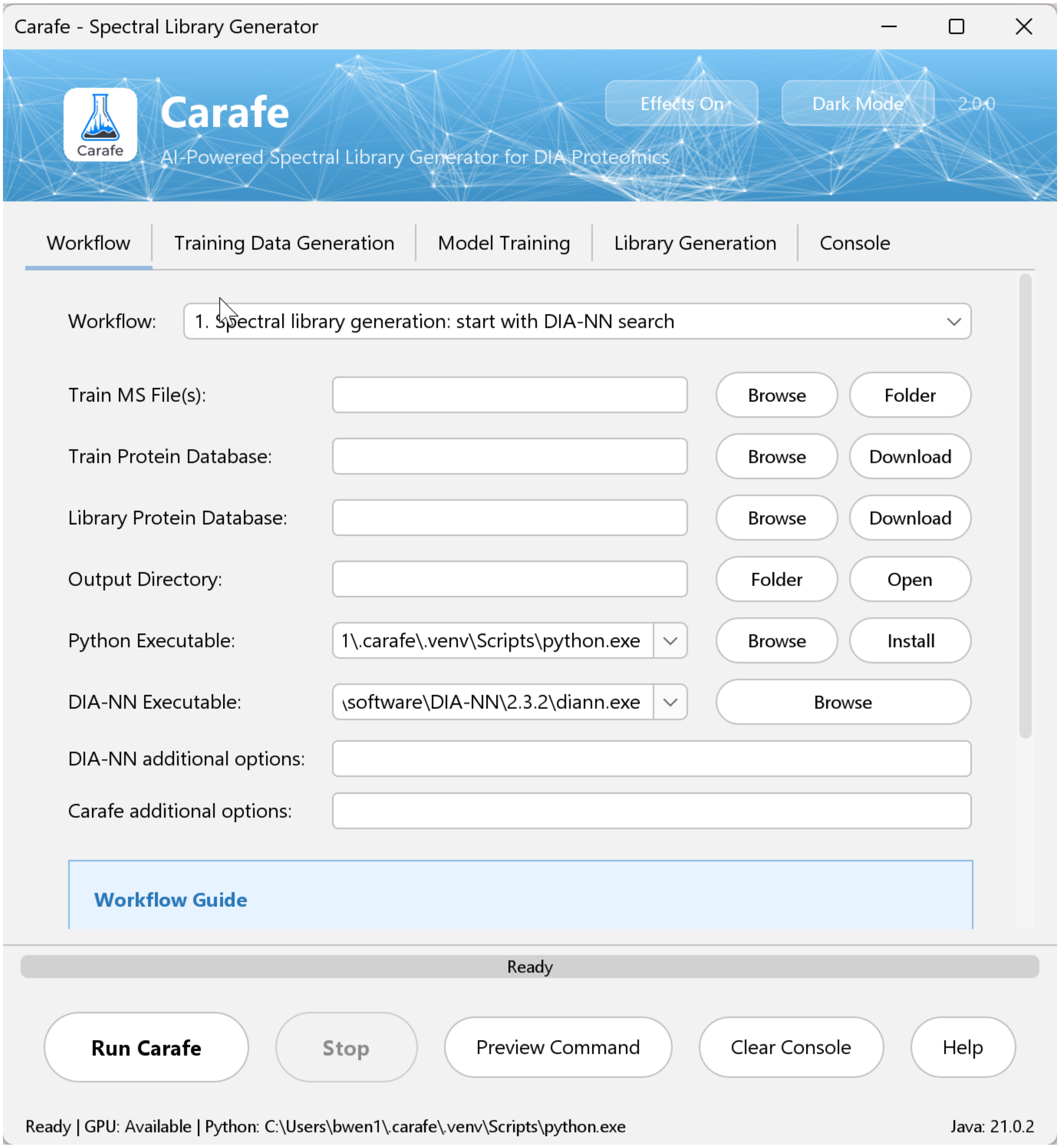


Supplementary Figure 1: **Carafe2 graphical user interface.** The Carafe2 GUI provides an intuitive interface for users to input parameters, select data files, and execute the spectral library generation process.

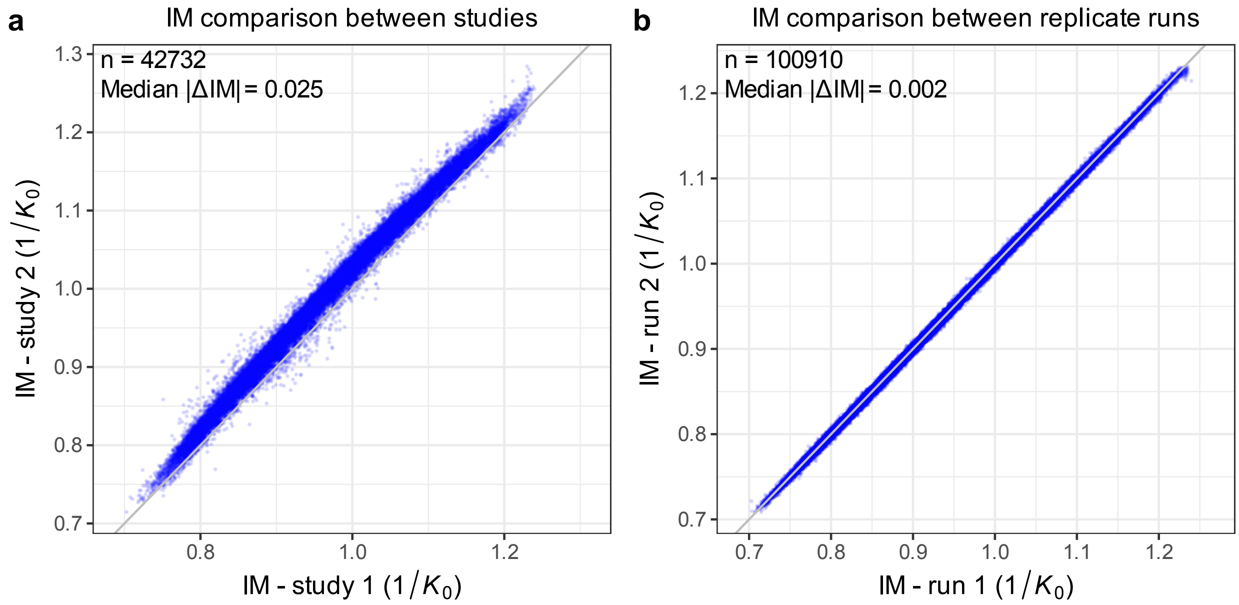


Supplementary Figure 2: **Comparison of ion mobility values between studies and replicate runs.** Scatter plots compare ion mobility values for matched peptide precursors detected in two MS runs. **(a)** Comparison between runs acquired in two independent studies (cross-study). **(b)** Comparison between two replicate runs acquired within the same study (within-study). Each point represents a matched precursor present in both runs; the gray diagonal indicates perfect agreement ($y = x$). Points are colored by ion mobility to aid visual tracking across the mobility range. Text annotations report the number of matched precursors ($n$) and the corresponding median absolute deviation shown in each panel.

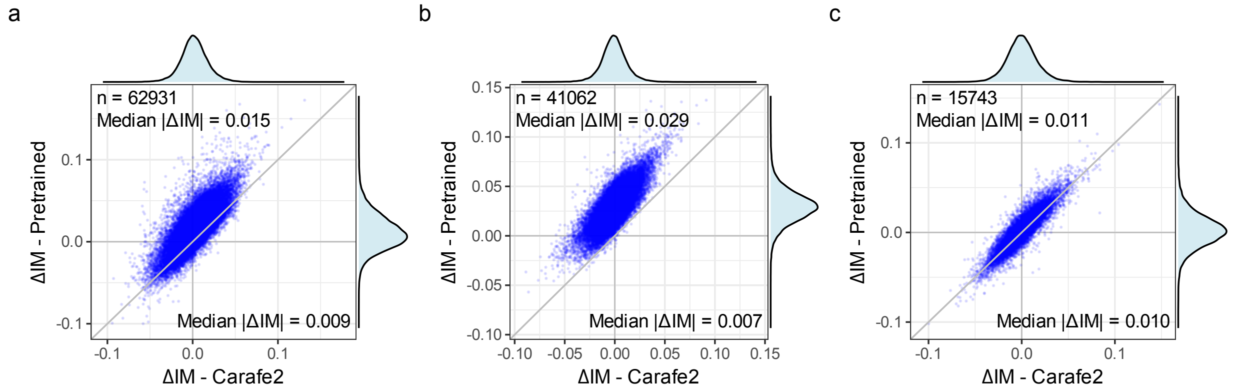


Supplementary Figure 3: **Comparison of ion mobility prediction between Carafe2 fine-tuned and pretrained models.** Scatter plots compare ion mobility prediction errors using Carafe2 fine-tuned models (x axis) and AlphaPeptDeep DDA data pretrained model (y axis) on the same three timsTOF DIA datasets used for Figure 3d–f, corresponding to panels **(a)**, **(b)**, and **(c)**, respectively. Text annotations report the number of matched precursors ($n$) and the corresponding median absolute prediction errors.

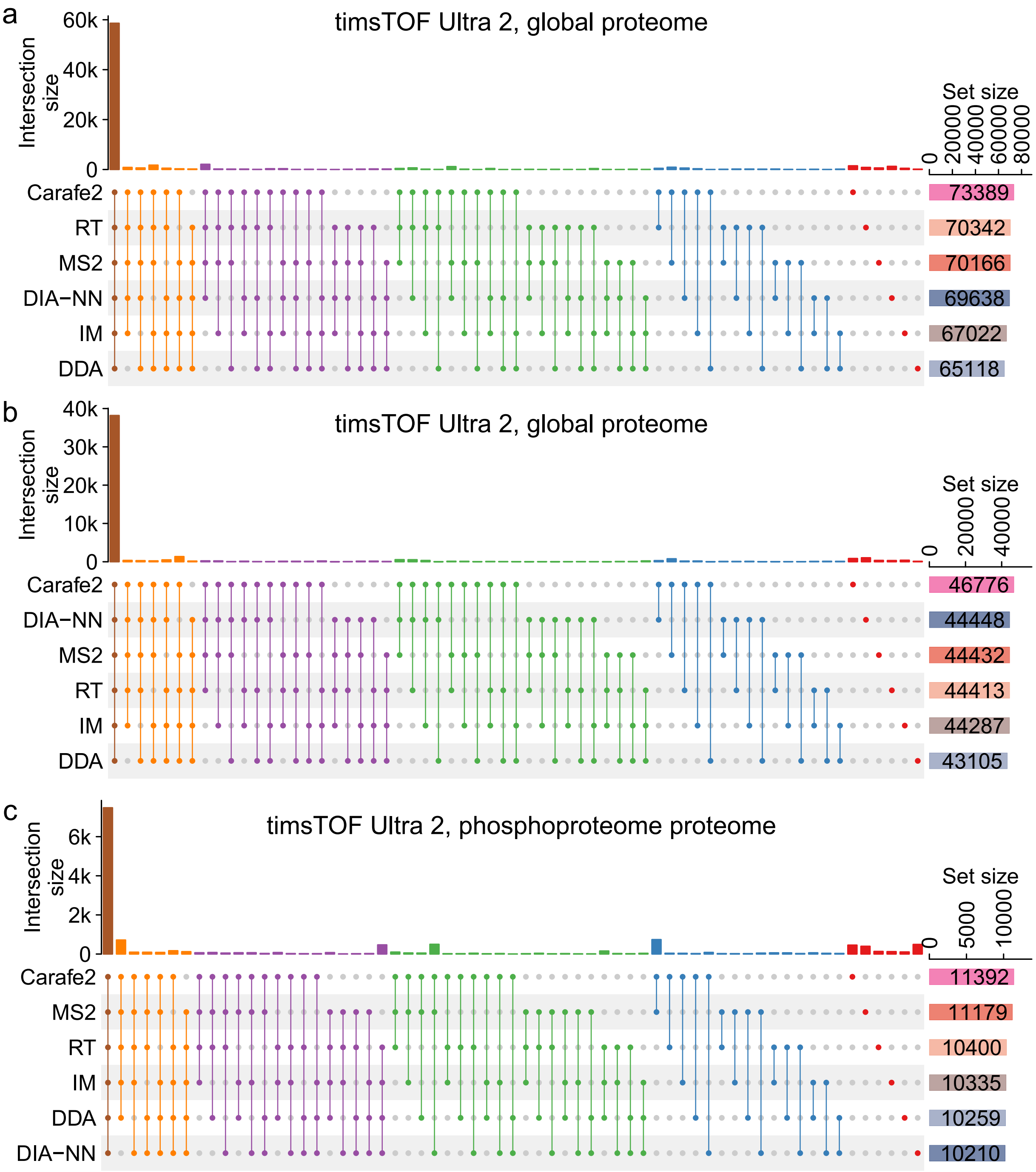


Supplementary Figure 4: **Upset plots showing the overlaps of peptide precursors detected using different *in silico* spectral libraries.** The upset plots correspond to the three datasets evaluated in Figure 4b–d of the main text: **(a)** a global proteome yeast dataset generated on a timsTOF Ultra 2 instrument, **(b)** a global proteome yeast dataset generated on a different timsTOF Ultra 2 instrument with a shorter LC gradient, and **(c)** a yeast phosphoproteome DIA dataset generated on a timsTOF Ultra 2 instrument. The dots representing combination sets are colored according to the number of sets included.

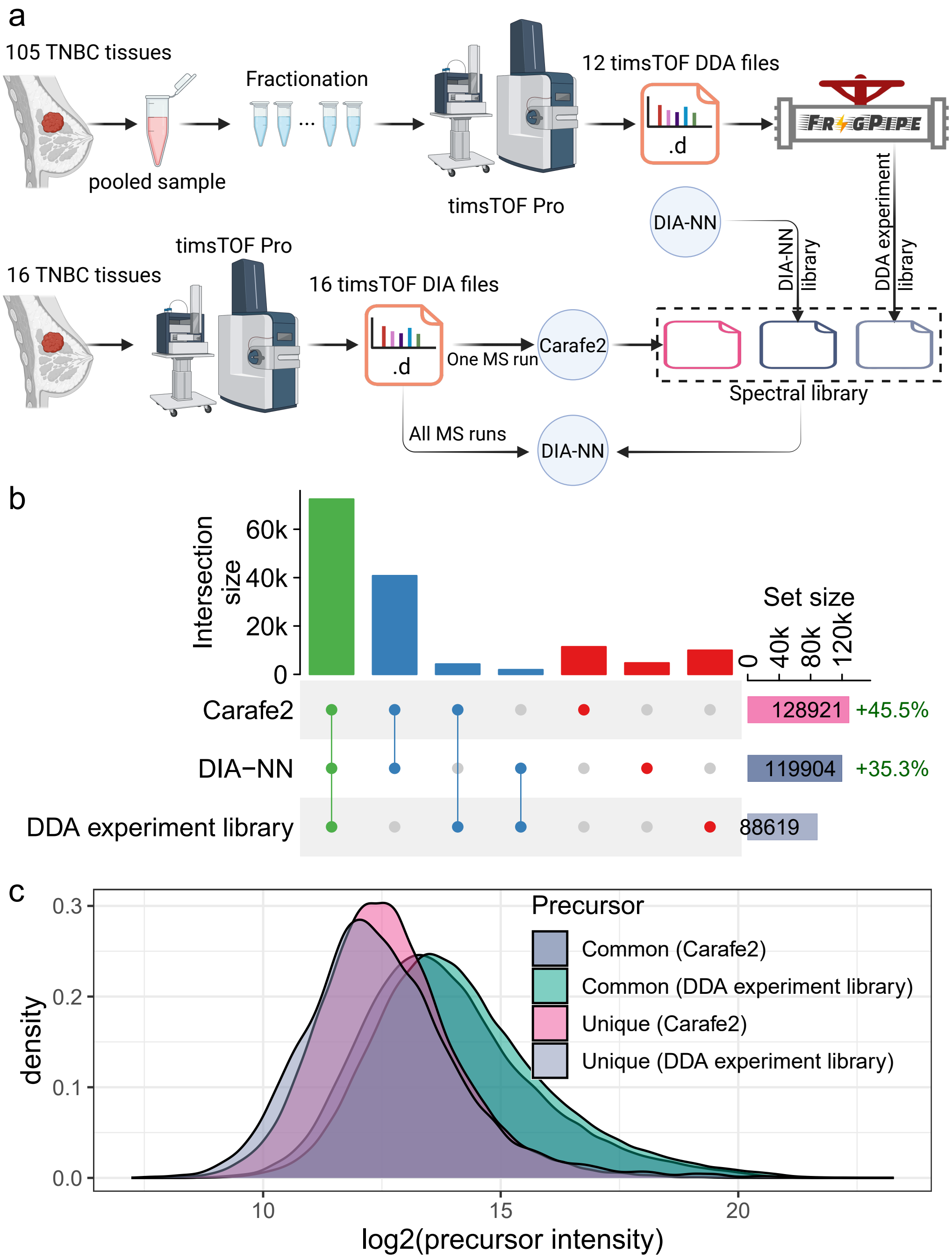


Supplementary Figure 5: **Comparison of *in silico* and DDA experimental spectral libraries. (a)** Comparing an experimental spectral library with *in silico* spectral libraries on a human tissue dataset. The DDA experimental spectral library was generated using FragPipe on 12 DDA timsTOF runs from the same study. The *in silico* spectral libraries were generated using DIA-NN and Carafe2, respectively. **(b)** An upset plot showing the number of precursors detected using different spectral libraries. **(c)** A precursor intensity density plot showing the distribution of precursor intensities for precursors detected between Carafe2 and DDA experiment library. Panel **(a)** was created using BioRender.
